# Peroxisome dynamics during HSV-1 life cycle in human neurons

**DOI:** 10.64898/2026.06.23.732381

**Authors:** Carolina Filipponi, Alessandro De Carli, Iwan Gane, Caterina Pignata, Elena Iacono, Fabio Filippini, Giulia Sciandrone, Domenico Favaro, Maribeth A Wesesky, Giulia Freer, Mauro Pistello, Leonardo D’Aiuto, Roberto Angelini, Michele Lai

## Abstract

HSV-1 is increasingly implicated in Alzheimer’s disease, yet the mechanisms by which it reshapes neuronal metabolism remain incompletely understood. Here, we demonstrate that HSV-1 co-opts peroxisomal biogenesis and lipid metabolic pathways to promote its replication across human neuronal models. In SH-SY5Y cells, infection triggers a marked expansion of the peroxisomal compartment and alters organelle morphology through upregulation of PGC-1α and PEX13/14/19. Pharmacological stimulation of peroxisome proliferation enhances viral production, whereas inhibition of PEX3–PEX19-dependent biogenesis almost completely suppresses infection. Lipidomic profiling reveals a selective increase in peroxisome-derived plasmalogens and sphingolipids, supporting a role for peroxisomes as a metabolic hub for viral envelopment. This remodeling is recapitulated in hiPSC-derived neurons and human brain organoids, where it is strictly dependent on productive replication and re-emerges upon viral reactivation, but not during latency. Collectively, these findings identify peroxisomes as essential replication-permissive organelles exploited by HSV-1 and suggest that recurrent virus-driven peroxisomal and ether-lipid reprogramming may contribute to neuronal vulnerability in neurodegenerative disease.

## Introduction

Over the past decade, Herpes Simplex Virus type 1 (HSV-1) has been increasingly associated with the progression of Alzheimer’s disease (AD)^1,2^. Experimental studies have shown that HSV-1 infection promotes the accumulation of phosphorylated tau and β-amyloid, two major hallmarks of AD, accompanied by microglial activation and sustained release of proinflammatory cytokines^3,4^. Despite this growing evidence linking HSV-1 to neurodegeneration, the impact of viral infection on neuronal metabolism and subcellular organelles remains poorly defined, particularly with regards to peroxisomes.

Peroxisomes play fundamental roles in neuronal physiology, including fatty acid β-oxidation, maintenance of redox homeostasis, and biosynthesis of ether lipids such as plasmalogens^5,6^. These lipids are critical structural components of neuronal membrane and myelin and are essential for synaptic function^7,8^. Altered peroxisomal dynamics and impaired plasmalogen metabolism have been repeatedly associated with AD progression and other neurodegenerative disorders, suggesting that peroxisomal dysfunction may contribute to neuronal vulnerability^9–11^. As reported by Semikasev et al., peroxisome abundance increases in the hippocampal and neocortical regions during milder disease stages. However, in more advanced stages, these areas exhibit a decline in peroxisome abundance^5,12^.

Recent evidence indicates that viruses can hijack peroxisomal pathways to support their replication and persistence^13^. For instance, human cytomegalovirus (HCMV), a member of the same *Herpesviridae* family as HSV-1, induces profound changes in peroxisome abundance, morphology, and lipid metabolism^14^. These findings raise the possibility that HSV-1 may similarly exploit peroxisomal function to favor infection and reactivation, with potential consequences for neuronal homeostasis.

In this study, we investigated how HSV-1 infection affects peroxisome dynamics and metabolism in neuronal systems. We first characterized the structural and transcriptional remodeling of peroxisomes during acute infection, focusing on peroxisomal abundance and plasmalogen biosynthesis. We started the analysis in neuroblastoma cells and then we moved to human induced pluripotent stem cell (hiPSC)-derived neurons and cortical organoids, which better mimic the cellular complexity of the human brain. Finally, using a latency-reactivation model, we explored how recurrent HSV-1 reactivation influences peroxisomal homeostasis. Together these approaches provide new insight into how HSV-1 reprogram peroxisomal metabolism, potentially linking viral persistence to neuronal dysfunctions.

## Methods

### Cell cultures, neuronal differentiation and brain organoids

Initial experiments were conducted in the SH-SY5Y neuroblastoma cell line due to its ease of handling and suitability for preliminary assays. However, given the tumor-derived nature of this line, human induced pluripotent stem cells (hiPSCs) were subsequently employed and differentiated into mature neurons and cortical organoids to more accurately recapitulate physiological neuronal mechanisms.

SH-SY5Y neuroblastoma cells were maintained in DMEM/F12 with 2 mM L-glutamine, non-essential amino acids and 10% fetal bovine serum (FBS, Sigma) at 37°C and 5% CO2. Medium was refreshed 2-3 times per week and tested for mycoplasma. For peroxisome modulation, cells were treated with 3 mM 4-phenylbutiryc acid (4-PBA) (MedChemExpress, USA, #Cat 1821-12-1) for 48 hours or NCC 55-0396 (MedChemExpress, USA, #Cat 357400) for 24 hours before HSV-1 inoculation and after the infection. Vehicle controls received DMSO. Drug toxicity was assessed by automated nuclear counts on a high-content imaging system, Operetta CLS.

hiPSCs were obtained from the National institute of Mental Health (NIMH) Center for Collaborative Studies of Mental Disorders-funded Rutgers University Cell and DNA Repository (RUCDR). hiPSCs were maintained under standard conditions (37°C, 5% CO2, 100% humidity). Neural progenitor cells (NPCs) were subsequently derived from hiPSCs as described by D’Aiuto et al.^15^. NPCs were expanded in NP expansion medium (Supplementary table 1). To plate NPCs for neuronal differentiation, cells were detached using Accutase (StemCell Technologies, USA) and gently centrifuged. Then, for neuronal differentiation, NPCs were seeded for 5 days in Incomplete Neurobasal Medium (Gibco, Thermo Fisher Scientific, USA) (Supplementary table 1) and half of the culture medium was replaced every other day. After 5 days, the cells were cultured in Complete Neurobasal medium for 6 weeks. The media were refreshed by replacing half of the culture medium every other day and the cell were differentiated for 4 weeks^16^.

hiPSCs were seeded at a density of 30.000 cells/well in low attachment 96-well plates, previously treated with Lipidure, a cell-repellent coating for surfaces, preventing the adhesion of 3D cell cultures. Cells were cultured in mTeSR1 (StemCell Technologies, USA) supplemented with rhFGF 10 μM and Rock Inhibitor 10 μM for 3 days. After Day 3 the medium was replaced with Essential 6 supplemented with dual-SMAD inhibitors, SB431542 (10 μM) and LDN193189 (100 nM) (Millipore Sigma, USA). On Day 8, the cells were moved to neuronal medium (Supplementary table 1) and then after 12 days, we substituted the media with CODMI Medium (Supplementary table 1). On Day 25 organoids were moved to CODMII Medium (Supplementary table 1) until day 42, when we replaced it with BrainPhys^TM^ Neuronal medium (StemCell Technologies, USA). The characterization of brain organoids differentiated using this protocol was performed by D’Aiuto and colleagues^16^.

### Generation of AGPS Knock Out SH-SY5Y

AGPS knockout SH-SY5Y cells were generated using CRISPR-Cas9-mediated gene editing. A crRNA targeting the first exons of AGPS (crRNA: GTACCAATGAGTGCAAAGCG) was annealed with Alt-R CRISPR-Cas9 tracrRNA in nuclease-free IDTE buffer to generate the guide RNA, according to the manufacturer’s instructions (Integrated DNA Technologies, Leuven, Belgium). The guide RNA was then complexed with Alt-R Cas9 protein to form ribonucleoprotein complexes. Approximately 150,000 SH-SY5Y cells were electroporated with the RNP complex and Alt-R Cas9 Electroporation Enhancer using the Neon MP5000 Electroporation System with Buffer R and Buffer E, following the manufacturer’s protocol (Invitrogen, Thermo Fisher Scientific, USA). Electroporation was performed using the following parameters: 1700 V, 20 ms, 1 pulse. Electroporated cells were cultured under standard growth conditions, selected using clonal selection, and AGPS depletion was assessed by western blot analysis and sequencing before downstream HSV-1 infection experiments.

### Viral propagation and infection

The HSV-1 strain McIntyre was purchased by ATCC (ATCC, CCL-539) and expanded at the University of Pisa. It was propagated on Vero E6 cells. Briefly, cells were infected at approximately 90% of confluence in DMEM 2% FBS until the appearance of cytopathic effects. The supernatant was then collected and centrifuged to eliminate the debris at 650 x g for 10 minutes. It was aliquoted and stored at -80°C. The KOS (ATCC, VR-1493) and *17syn+* (obtained by Prof. David D. Bloom, University of Florida) strains were expanded at the University of Pittsburgh on Vero cells. Viral titers were calculated by plaque assay and expressed as plaque-forming units (PFU)/mL.

The ZIKV virus strain MR-766 was purchased by ATCC (ATCC, VR-1838) and expanded at the University of Pisa. It was propagated on Vero E6 cells.

For SH-SY5Y infections, HSV-1 McIntyre was used at MOI 1 (kinetics starting from 4 to 48 hpi) or MOI 0.1 (drug-modulation experiments). Cells were inoculated in serum-free medium for 1 hour, washed, and maintained in complete medium. SH-SY5Y cells were infected with ZIKV at an MOI of 2 for 1 hour and 3 minutes and then cultured for 48 hours to maximize infection.

hiPSC-derived neurons were infected with HSV-1 17syn+ and KOS at an MOI of 1 for 6 to 24 hours, with 1 hour adsorption in DMEM/F12 followed by maintenance in Neurobasal medium.

Organoids were infected with HSV-1 17syn+ and KOS using 3000 PFU per organoid in a small volume of Neurobasal medium. For latency, organoids were pretreated with 30 μM (E)-5-(2-bromovinyl)-2’-deoxyuridine (5BVdU) and 125 U/μL Interferon-α (IFN-α), infected with 1500 PFU HSV-1 strain KOS, and maintained with or without antiviral/IFN-α before switching to BrainPhys. Reactivation was induced by 5 mM sodium butyrate (NaB) or 20 μM LY294002 for 5 days. This latency protocol was validated in Prof. Leonardo D’Aiuto’s Laboratory^16^.

### Plaque assay

Plaque assays were carried out using VeroE6 cultured in DMEM. Cells were infected using serial dilutions of HSV-1 for 1 hour at 37°C in 5% CO2. Following adsorption, the inoculum was removed, and cells were overlaid with culture medium containing 0.75% methylcellulose supplemented with 2% FBS. After 48 hours of incubation, cells were fixed in paraformaldehyde overnight and subsequently stained with crystal violet.

### RNA extraction and RT-qPCR

The RNA extraction was performed from the cell pellet to analyze the modulation of gene expression after HSV-1 infection, including genes involved in peroxisomal biogenesis, such as PEX13, PEX14 and PEX19, and genes involved in the plasmalogen biosynthetic pathway, GNPAT, FAR1, AGPS, EPT1. SH-SY5Y cells were seeded and infected with HSV-1 strain Mc Intyre the following day. Infection was terminated at 4, 8, 12, 16, 24 and 48-hours post-infection. Total RNA was extracted using QIAzol Lysis Reagent (QUIAGEN, Hilden, Germany) according to the manufacturer’s protocol. RNA concentration and purity were assessed using a NanoDrop spectrophotometer (Thermo Fisher Scientific, MA, USA). The RNA extraction of differentiated neurons was performed using the QIAgen RNeasy Plus Mini Kit (QUIAGEN, Hilden, Germany) to improve the quality of the extracted RNA with effective gDNA removal using gDNA Eliminator columns. The isolation was performed according to the manufacturer’s instructions provided with the kit.

Gene expression of PEX13, PEX14, PEX19, GNPAT, FAR1, AGPS, and EPT1 was measured by SYBR Green RT-qPCR with a one-step protocol, using β-actin (SH-SY5Y) and HPRT (neurons) as reference genes. Gene expression was calculated using the ΔΔCt method.

The primer used for RT-qPCR are resumed in Supplementary table 2.

### Immunofluorescence

#### Immunofluorescence of SH-SY5Y and high content confocal imaging

SH-SY5Y cells were seeded in 96-well PhenoPlates (Revvity, Waltham, MA, USA). SH-SY5Y cells were fixed in 4% paraformaldehyde (PFA) for 20 minutes. Samples were blocked for 1 hour at room temperature in blocking buffer (PBS containing 0.3% Triton X-100, 0.1% goat serum, and 3% human serum) and incubated at room temperature for 2 hours with the following primary antibodies: mouse monoclonal anti-HSV-1 gD (Invitrogen, Cat# G610D, 1:200), rabbit anti-PEX14 (Abcam, Cat# ab183885 1:400), mouse anti-flavivirus NS1 (Abcam, Cat#ab214337, 1:400). Then, the cells were washed with PBS and incubated with fluorophore-conjugated secondary antibodies for 1 hour at room temperature. The secondary antibodies used were the following: goat anti-rabbit Alexa Fluor 568 (Invitrogen, Cat# A11011, 1:1000) and goat anti-mouse Alexa Fluor 488 (Invitrogen, Cat# A11029, 1:1000). Nuclei were stained with DAPI (Sigma Aldrich, St. Louis, MO, USA, 1:1000) and cytoplasm was labelled using phalloidin 647 (Abcam, Cat# ab176759, 1:2000). Samples were imaged using Operetta CLS High-Content Imaging System (Revvity, Waltham, MA, USA). In the data acquired with Operetta CLS High-content confocal microscopy, each data point of the graph represents a mean obtained after the analysis of at least 8×10^3^ cells.

For image analysis we used Harmony 4.6 Software (Revvity, Waltham, MA, USA) following the building blocks: Find nuclei > Find cytoplasm > Calculate intensity properties (HSV-1 gD – Alexa 488) > Select population: Infected cells > Find spot (PEX14 – Alexa 568) in infected and uninfected populations > Calculate Morphology properties of spots. Around 150 fields were analyzed per well, using a 40x water objective. PEX14 spots were identified and filtered based on fluorescence intensity, and the results were expressed as peroxisome number arbitrary unit (AU) per cell.

#### Immunofluorescence of hiPSCs-induced neurons and brain organoids

Organoids were fixed in 4% PFA overnight at 4°C, washed three times in PBS, and cryoprotected by incubation in a 30% sucrose solution overnight. The organoids were then transferred into cryomolds (Tissue-Tek Cryo Mold Intermediate). Samples were embedded in OCT medium and frozen at -22°C. Cryosection of 10 μm were obtained using a cryostat (Micron HM350, Thermo Fisher Scientific, Pittsburgh, PA, USA) and stored at -80°C until use. Prior to staining, sections were air-dried, post-fixed with 4% PFA for 20 minutes, and permeabilized/blocked in 10% normal goat serum with 0.02% Triton X-100 for 1 hour at room temperature. Sections were then incubated overnight at 4°C with the following primary antibodies: mouse anti-HSV1 ICP4 (Abcam, Cat# ab6514, 1:200), rabbit anti PEX14 (Abcam, Cat# ab183885 1:400). The next day, the samples were stained with fluorophore-conjugated secondary antibodies including Alexa Fluor 488 goat anti-mouse (Thermo Fisher Scientific, Cat# A-10680, 1:300) and Alexa Fluor 594 goat anti-rabbit (Thermo Fisher Scientific, Cat# A-11012, 1:300). Nuclei were stained using Hoechst dye (1:1000) for 5 minutes. The 2D neuronal cultures were fixed with 4% PFA and then stained as described for brain organoids, using a permeabilization/blocking solution of 10% normal goat serum with 0.2% Triton X-100.

Fluorescence images were acquired using a Leica CTR5500 microscope for representative images, and a Nikon A1 point-scanning confocal microscope for quantitative analysis with a 63x magnification objective. Image processing and quantification were performed using Fiji, measuring the fluorescence intensity and the number of PEX14 spots. Correct total fluorescence intensity (CTCF) was obtained normalizing for background using the following equation: CTCF = Measured fluorescence – (Area of selected region x Mean of the background). PEX14 spots were identified and filtered based on fluorescence intensity, and the results were expressed as peroxisome number arbitrary unit (AU) per cell. Each data point derived from the analysis of samples acquired using classical confocal microscopy stands for the mean values obtained from the analysis of at least 30 cells.

### Western Blot analysis

Cells were washed with PBS and lysed in RIPA Buffer supplemented with protease inhibitor cocktail (Thermo Fisher Scientific, USA). Following a 2-hour incubation at 4°C, lysates were centrifuged for 10 minutes at 4°C at 13.000xg. Polyacrylamide gels were prepared using SureCast Acrylamide Solution (40%) (Thermo Fisher Scientific, USA). For optimal separation of the interest proteins, we prepared a 12% resolving gel and a 4% stacking gel. Protein samples were denatured at 70°C for 10 minutes in 5X Loading Buffer (25 0nM Tris-HCl pH 6.8, 10% SDS, 30% glycerol, 0.05% bromophenol blue, 0.05% 2-mercaptoethanol). Electrophoresis was performed in running buffer (25 mM Tris base, 192 mM glycine, 0.1% SDS, pH 8.3) at 200V for 1 hour. Proteins were then transferred into nitrocellulose membrane 0.22 μm using transfer buffer (25 mM Tris base, 190 mM glycine, 20% methanol) at 350 mA for 1 hour. Membranes were blocked for 1 hour at room temperature in 5% nonfat dry milk in PBS Tween 0.1%. Primary antibodies used included mouse anti-HSV-1 glycoprotein D (Santa Cruz, Cat# sc-21719, 1:1000) and rabbit anti-GAPDH (Invitrogen, Cat# MA5-15738, 1:2000). Primary antibodies were incubated for overnight at 4°C. After washings, the membranes were incubated with HPRO-conjugated anti-mouse IgG (Sigma, Cat#A9044, 1:20000) and anti-rabbit IgG (Sigma, Cat#A0545, 1:20000) for 1 hour at room temperature.

### Lipid extraction

#### Protein quantification

SH-SY5Y cells were infected with HSV-1 strain Mc Intyre at an MOI of 1. Cells were then pelleted, and the pellet was heat-inactivated at 70°C for 10 minutes. The samples were resuspended in 155mM ammonium bicarbonate (AB, NH4HCO3), and tip sonicated to homogenize the samples before use. To ensure equal loading of the samples, protein concentrations were quantified using the BCA Protein Assay Kit (Thermo Fisher Scientific, Pittsburgh, PA, USA, Cat# A55865).

#### Lipid extraction

To estimate the lipid content of infected cells, we performed a lipid extraction using a modified Bligh & Dyer chloroform/methanol protocol, adapted for small cell samples^17^. Briefly, cell lysates containing 100 μg of total protein were adjusted to a final volume of 400 μL with AB solution. A mixture of internal standards (300 pmoles of PE 12:0/12:0, 200 pmoles of PS 14:0/14:0, 200 pmoles of Cer 18:1;2/12:0;0, 200 pmoles of PC 12:0/12:0, 200 pmoles of SM 18:1;2/ 12:0;0) was then added to each sample to control extraction efficiency and provide quantification. Lipid extraction was initiated by adding 1 mL of methanol containing 2% butylated hydroxytoluene (BHT) as an antioxidant and 500 μL of chloroform. The samples were vortexed for 1 minute and incubated at room temperature for 30 minutes. Subsequently, 500 μL of AB solution and 1.5 mL of chloroform were added, followed by 1 min vortexing, resulting in the formation of a biphasic system.

Samples were then centrifuged at 1.000 x g for 10 minutes to facilitate phase separation. The lower organic phase was carefully collected into a new glass tube using a glass Pasteur pipette.

The organic phases were dried under a gentle stream of nitrogen at 45°C to concentrate the lipid extract. The dried samples were resuspended in 100 μL of chloroform, and the lipid extracts were then stored at -20°C until further analysis.

### Electrospray ionization mass spectrometry analysis

Lipid extracts were analyzed by direct-infusion electrospray ionization (ESI) mass spectrometry using a TriVersa NanoMate (Advion, NY, USA) as the nanospray interface. A spray voltage was applied to generate ionization, and data were acquired in both positive and negative ion modes, according to the lipid class under investigation.

The mass spectrometer used was an LTQ-Orbitrap hybrid system (Thermo Fisher Scientific, USA), in which ions are separated based on their mass-to-charge ratio (m/z) through high-resolution orbital trapping. Collision-induced dissociation (CID) was performed for structural characterization and confirmation of lipid species.

For negative ion mode analysis, 5 μL of lipid extract were combined with 8 μL of 0.2% diethylamine (DEA) in chloroform/methanol (1:5, v/v). For positive ion mode, 5 μL of lipid extract were mixed with 8 μL of 13.3 mM ammonium acetate (NH4Ac) in 2-propanol. The prepared mixtures were loaded into the wells of a 96-well NanoMate plate and directly infused into the mass spectrometer for lipid analysis (i.e., shotgun lipidomics)^18^.

Our analytical method enabled the identification of different species, including pPE. In assigning pPE, we followed the conventional assumption on the presence of vinyl ether bond at the sn-1 position of the fatty alcohol chain. This assumption is well supported by literature, which consistently reports that in the brain pPE species almost exclusively predominates over PEo^19,20^. Consequently, when a double bond is detected in ether-linked chain, it can be reasonably attributed to a vinyl ether bond.

### Statistical analysis

Data was reported as mean ± SD from independent experiments. The number of technical replicates per biological condition was 2 or 3, depending on the specific analysis. Statistical significance was determined using one-way analysis of variance (ANOVA), Multiple comparison Tuckey’s testo Multiple comparison Dunnett’s test, to compare more than two groups, and an unpaired two-tailed Student’s *t*-test to compare two groups. Differences were considered statistically significant at p < 0.05 (*), p < 0.01 (**), p < 0.001 (***), p < 0.0001 (****) with α=0,5.

All statistical analyses were performed using GraphPad Prism (GraphPad Software, San Diego, CA, USA).

## Results

### Peroxisomal rearrangement occurs after HSV-1 infection in SH-SY5Y cells

Since peroxisomes play an important role in viral infections, we aimed to analyze peroxisome biogenesis and functions during the progression of HSV-1 infection. Given the complexity and tight regulation of peroxisome biogenesis, we assessed PGC-1α expression, the primary regulator of PEX11β, a major activator of peroxisomal biogenesis^21,22^, along with three PEX genes involved in matrix protein import, PEX13 and PEX14, and in PMPs import, PEX19^23^. The analyses were performed in the presence or absence of HSV-1 infection. The neuroblastoma cell line SH-SY5Y was employed for an initial screening, in which cells were infected with HSV-1 (strain McIntyre) at an MOI of 1 and analyzed at 4,6,16,24 and 48 hours post infection (hpi), as schematically illustrated in Figure 1a.

**Figure 1.**
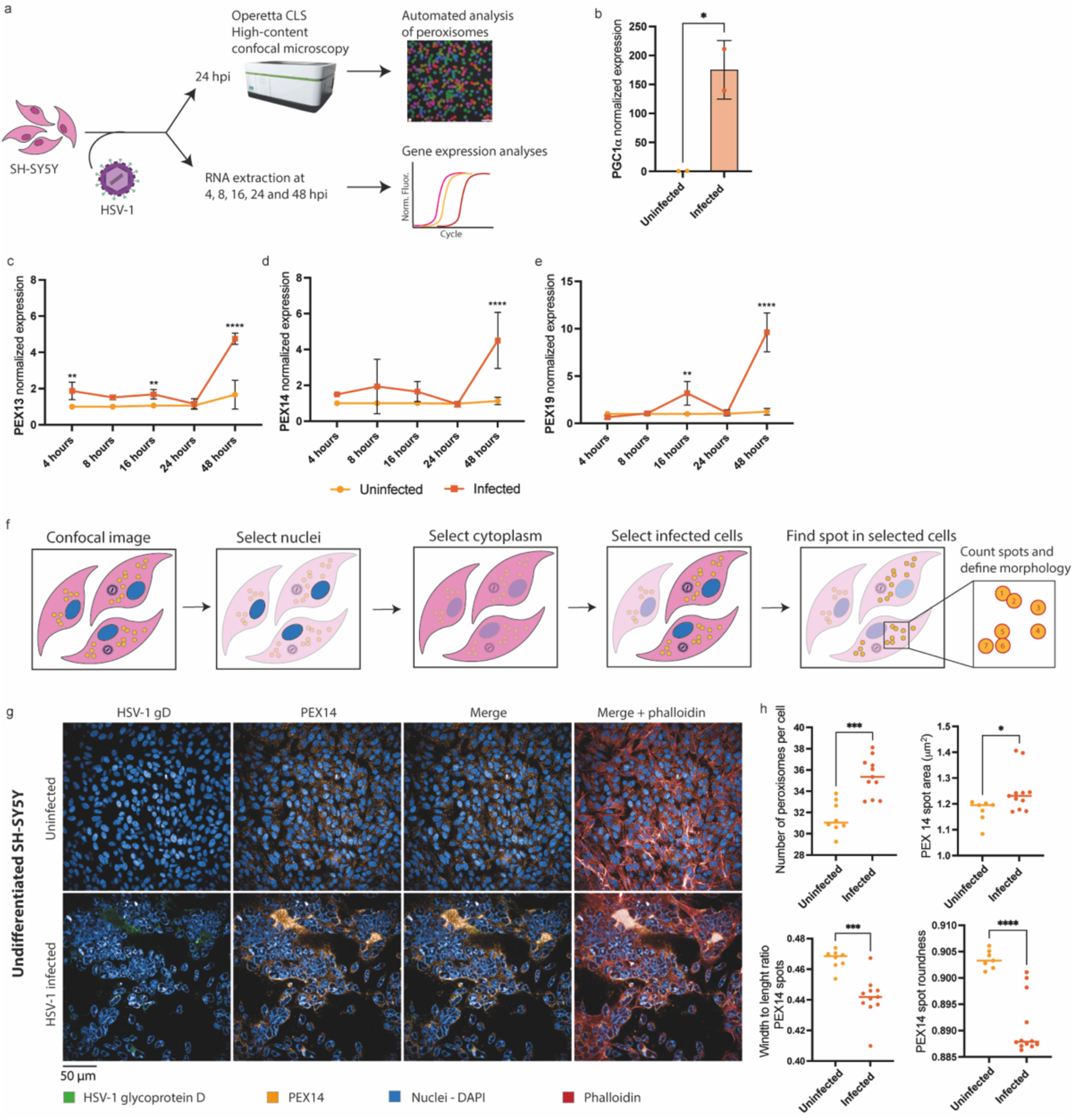
HSV-1 infection modulates peroxisome. a. Overview of the experimental workflow. Briefly, SH-SY5Y cells were infected or not with HSV-1 strain McIntyre at an MOI of 1. The infected cells and control samples were stained and analyzed using high content confocal microscopy. The analysis of peroxisome number and morphology was obtained using the highly automated Harmony Software. Additionally, RNA extraction was performed, and RT-qPCR was used for PEX gene expression at diQerent time points (4, 8, 16, 24, 48 hpi). b. Gene expression analysis of a key activator of peroxisomal biogenesis PGC1α. c, d, e. Gene expression analyses of PEX13, PEX14 and PEX19 at diQerent time point of infection. Data were analysed with Student’s t-test comparing uninfected and infected conditions within the same time point. Results are expressed as mean ± SD of six independent replicates. f. Schematic representation of image analysis using the highly automated software, Harmony. g. Panel of representative confocal microscopy images. Peroxisomes were labeled with PEX14 (orange), HSV-1 was visualized using HSV-1 gD (green), nuclei with DAPI (blue) and the cytoskeleton with phalloidin (red). The images were acquired using Operetta CLS high content confocal microscopy with a 40x magnification. h. Analyses of peroxisome number per cell and morphology. Each data point represents the mean value obtained analysing > 8×10^3^ cells. Results are expressed as mean ± SD of independent replicates. Data were analysed with Student’s t-test comparing uninfected and infected conditions (* p < 0.05, ** p < 0.01, *** p < 0.001; **** p < 0.0001, α=0.5, N=>8×10^3^/sample).

Quantitative RT-PCR analyses revealed a 100-fold upregulation in the expression of PGC1α at 24 hpi (Figure 1b). Next, we assessed the temporal expression profile of the PEX genes. PEX13 expression was increased approximately 2-fold at 4 hpi, 1.5-fold at 16 hpi, and 5-fold at 48 hpi (Figure 1c). PEX14 expression was upregulated 4-fold at late infection stages (48 hpi, Figure 1d), while PEX19 showed a 2-fold increase at 16 hpi, reaching a 10-fold increase at 48 hpi (Figure 1e).

Since peroxisome biogenesis genes were upregulated following HSV-1 infection, we next analyzed peroxisome morphology and abundance. Peroxisomes were labeled using PEX14 immunostaining and visualized 24 hpi using high-content confocal microscopy, as depicted in Figure 1f. We observed a 14% increase in the total number of peroxisomes per cell (AU) in infected cells. Intriguingly, peroxisomes appeared enlarged and more elongated exhibiting a reduced width-to-length ratio compared with infected controls (Figure 1g-h).

Gene expression and immunofluorescence analyses together indicate that HSV-1 infection drives extensive peroxisomal remodeling. The coordinated upregulation of biogenesis genes and the accompanying morphological alterations suggest that peroxisomes actively participate in HSV-1 infection, either by contributing to antiviral defense or by being hijacked to sustain viral replication. It remains unclear whether the observed morphological changes reflect virus-driven remodeling of the peroxisomal compartment or instead represent physiological peroxisomal elongation associated with the normal *growth-and-fission* cycle of biogenesis.

### HSV-1 directly uses peroxisomes for its replication

Since peroxisomes were modulated during HSV-1 infection in SH-SY5Y cells, we next examined whether they contribute to an antiviral host immune response or instead have a proviral activity, facilitating viral replication. The experimental workflow is summarized in Figure 2a. SH-SY5Y cells were pretreated with 4-phenylbutyrate (4-PBA), a chemical enhancer of peroxisomal proliferation^14^, or with NCC 55-0396, a selective inhibitor that blocks PEX3–PEX19 interaction and thereby suppresses peroxisome biogenesis^24^, 48 or 24 hours prior to HSV-1 infection, respectively. Following pretreatment, cells were infected with HSV-1 and analyzed after 24 hours by high-content confocal microscopy and viral yield assays. Drug toxicities are shown in Supplementary figure 1 and representative confocal microscopy images of the different conditions are shown in Figure 2b. Cells treated with 4-PBA showed a robust induction of peroxisome formation, increasing the mean number of peroxisomes per cell from 41.5 to 60.4 arbitrary fluorescence units (AU), corresponding to an approximate 50% increase compared with vehicle-treated controls (Figure 2c). Morphological analysis further revealed peroxisomal changes upon 4-PBA treatment (Figure 2d) that were comparable to those observed during HSV-1 infection alone (Figure 1h), suggesting that the observed morphological alterations likely reflect normal peroxisomal proliferation rather than a virus-specific pathological mechanism.

**Figure 2.**
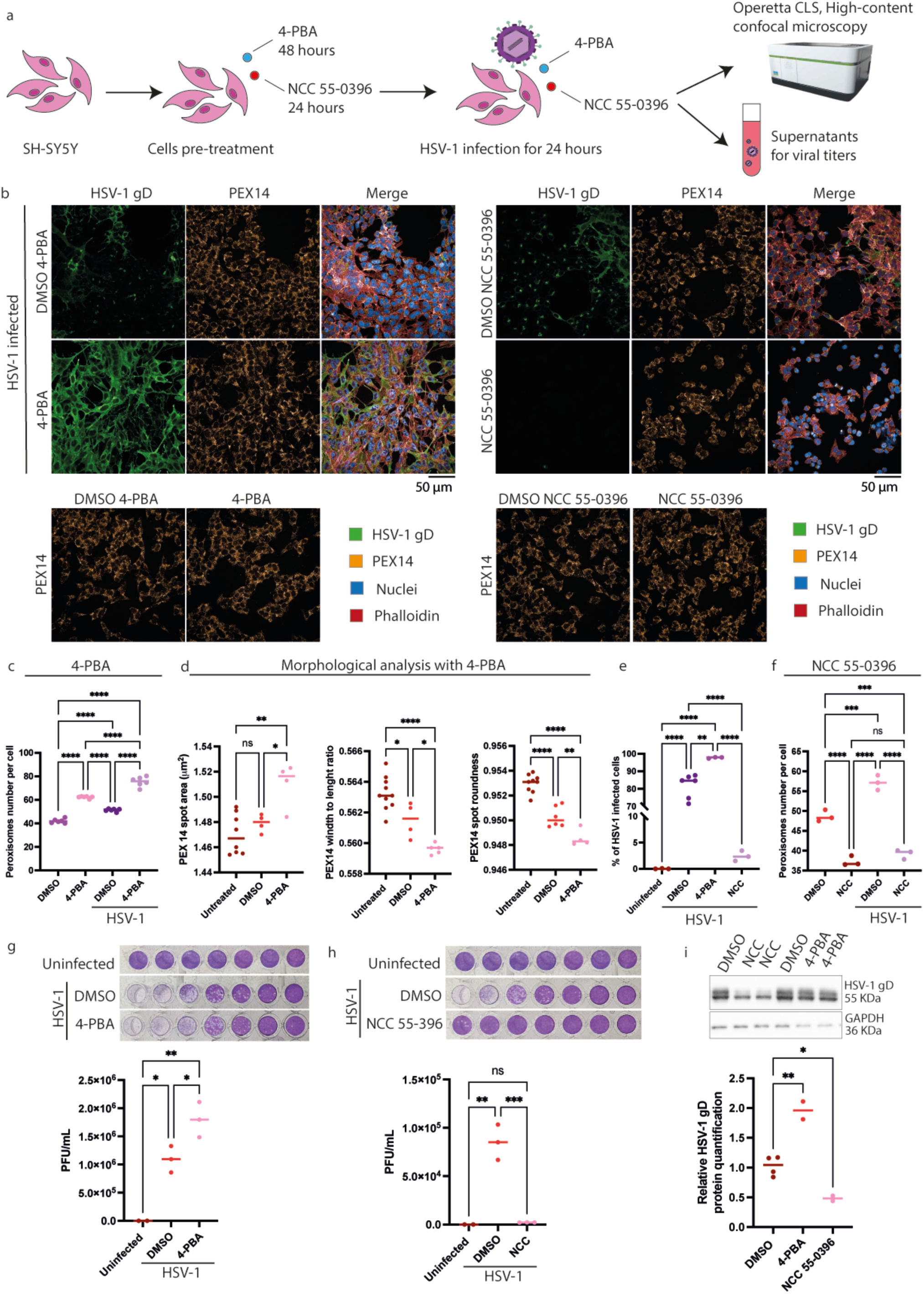
4-PBA and NCC 55-0396 modulates HSV-1 infection. a. Brief description of the workflow. SH-SY5Y cells were pretreated or not with 4-PBA for 48 hours and with NCC 55-0936 for 24 hours. Then, the cells were infected with HSV-1 strain McIntyre at an MOI of 0.1 and analyzed using high content confocal microscopy, viral yield and western blot analysis. b. Panel of confocal microscopy images representing the diQerent conditions. Peroxisomes were labelled using PEX14 (orange), HSV-1 using HSV-1 gD (green), nuclei using DAPI (blue), and cytoskeleton using phalloidin (red). Images were acquired using a 40x magnification. Each data point represents the mean value obtained analysing > 8×10^3^ cells. c. Analysis of number of peroxisomes per cell after the treatment with 4-PBA and DMSO. d. Morphological analysis of peroxisomes expressed as spot area, width to length ratio, and roundness after the treatment with 4-PBA. e. Analysis of the percentage of infected cells 4-PBA and NCC 55-0396 treated compared to the vehicle treated control. f. Analyses of peroxisomes per cell after the treatment with NCC 55-0396 and DMSO as vehicle control. g. Viral titer of the cell treated with 4-PBA and DMSO expressed as PFU/mL. h. Viral titer of the cell treated with NCC 55-0396 and DMSO expressed as PFU/mL. i. Western blot analysis on HSV-1 gD content following treatments. GAPDH was used as loading control. Results are expressed as mean ± SD of independent replicates. Data were analysed with One-Way ANOVA (* p < 0.05, ** p < 0.01, *** p < 0.001; **** p < 0.0001)

Interestingly, the infection rate of 4-PBA treated cells was approximately 50% higher compared to the vehicle-treated controls (Figure 2e), suggesting that increased peroxisomal abundance is a proviral factor promoting HSV-1 replication rather than being involved in antiviral host response. Moreover, the effects of HSV-1 infection and 4-PBA treatment appeared additive, as peroxisome numbers were further elevated when both stimuli were combined (Figure 2e).

To further confirm the direct involvement of peroxisomes, we used the compound NCC 55-0396, which inhibits PEX3-PEX19 interaction and thereby blocks peroxisome biogenesis^24^. Conversely, treatment with NCC 55-0396 reduced peroxisome spots (AU) from 48.3 to 36.7 per cell (Figure 2f), and markedly suppressed infection, decreaseing the HSV-1-positive cells from ∼80% to < 5% (Figure 2e).

Viral yield assays and western blot analysis confirmed these findings. Cells treated with 4-PBA exhibited approximately two-fold higher viral titers than DMSO-treated control (Figure 2g), whereas NCC 55-0396 treatment reduces viral replication (Figure 2h). Western blot analysis confirmed an accumulation of HSV-1 gD protein in the 4-PBA treated cells and a reduction where NCC 55-0396 was administered (Figure 2i).

Together, these findings corroborate the hypothesis that peroxisome biogenesis is required for efficient HSV-1 replication. Enhancing peroxisomal proliferation promotes viral production, whilst inhibition of their biogenesis severely compromises viral replication. This evidence was further confirmed by analyzing PEX14 spots in uninfected cells from the same wells where HSV-1 infection occurred. We found that the number of PEX14+ spots in uninfected cells are significantly lower than in surrounding infected cells, ruling out soluble antiviral factors as the cause of increased peroxisomes and instead implicating HSV-1 replication itself (Supplementary figure 2).

To assess whether peroxisomal alterations represent a general feature of enveloped viruses or are specific to HSV-1 and herpesviruses, we analyzed cells upon infection with ZIKV, an enveloped RNA virus that replicates in the endoplasmic reticulum/cytoplasm. Quantification of peroxisomes by automated confocal analysis revealed a 20% reduction in the average number of peroxisomes (AU) per cell following ZIKV infection (Supplementary figure 3b, c). Interestingly, pretreatment with 4-PBA led to modest 8% decrease in ZIKV infection rate and a marked 95% reduction in viral titers compared to vehicle-treated controls (Supplementary figure 3b, c). These results indicate that, unlike HSV-1, ZIKV replication is negatively affected by increased peroxisome abundance, supporting the existence of virus-specific strategies in peroxisome-virus interactions.

### HSV-1 infection alters plasmalogen biosynthesis and lipid composition in SH-SY5Y cells

Our findings confirm those observed by Beltran and colleagues^14^, in which HCMV also increases peroxisome numbers in the late stage of infection. To further confirm this mechanism, we hypothesized that HSV-1 might also rely on peroxisomes to produce plasmalogens, a lipid class that have to be incorporated into the virions. To do so, SH-SY5Y cells were infected with HSV-1 at MOI 1 and the expression of key peroxisomal enzymes involved in plasmalogen biosynthesis was analyzed. Specifically, we quantified GNPAT and AGPS, which catalyze the first two steps of ether lipid formation in peroxisomes, as well as the rate-limiting enzyme in the pathway FAR1, which supplies fatty alcohols for ether lipid production, and EPT1, a downstream enzyme that maintains plasmalogen homeostasis, as schematically illustrated in Figure 3a.

**Figure 3.**
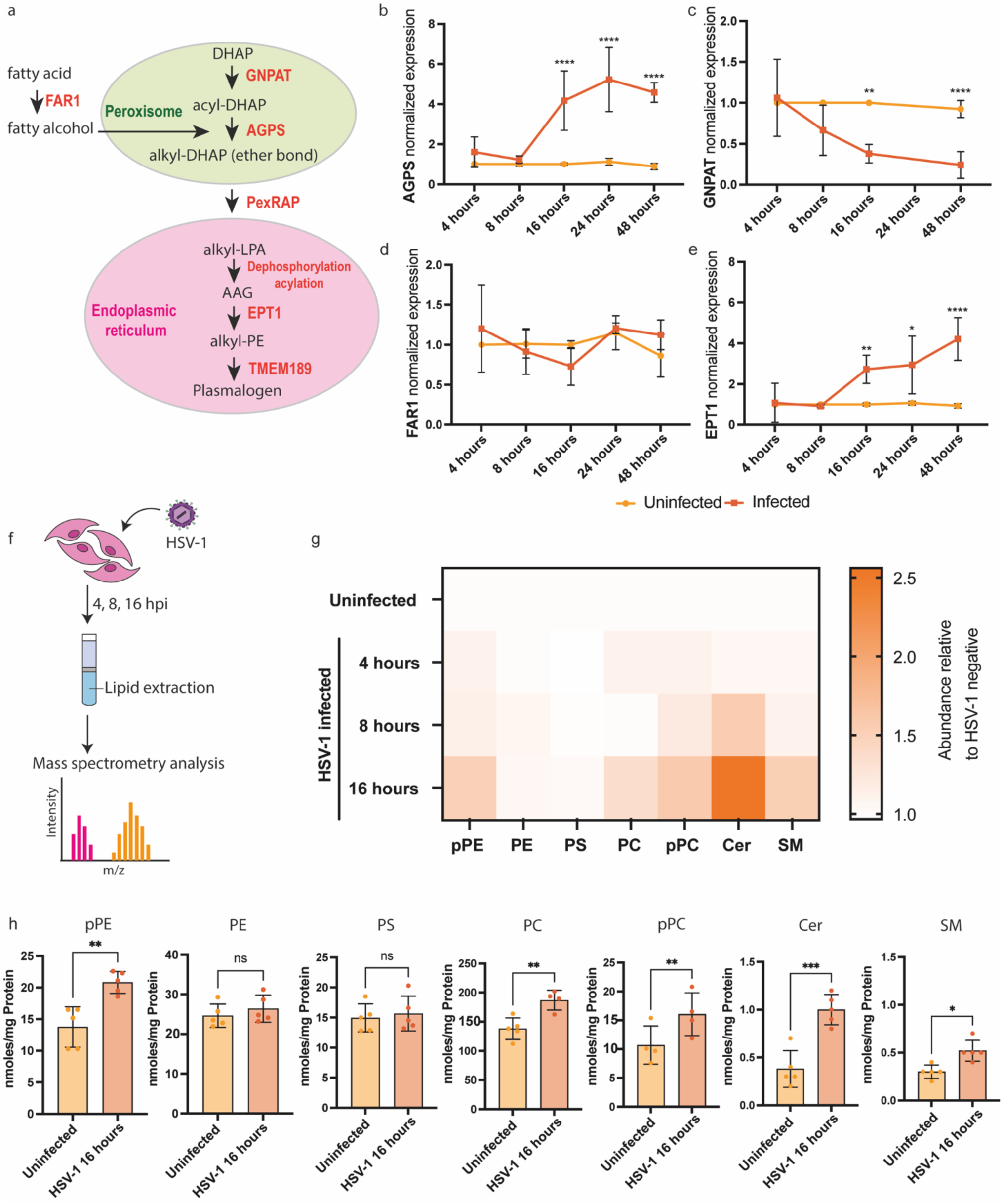
Plasmalogen metabolism increases after HSV-1 infection. a. Schematic representation of peroxisomal biosynthetic pathway. b, c, d, e. Gene expression analyses of four genes involved in diQerent steps of plasmalogen production. The gene expression analyses were performed at diQerent time point after infection (4, 8, 16, 24, 48 hpi). f. Overview of the lipid analysis workflow. Briefly, the SH-SY5Y were seeded and infected at 1 MOI with HSV-1 strain McIntyre. The cells were then harvested after 4, 8, 16 hours, a lipid extraction was performed, and mass spectrometry analysis was run. g. Heatmap representing the abundance relative to HSV-1 negative samples of the diQerent lipid species analyzed at diQerent time points. h. Absolute quantification expressed in nmol/mg of protein of the diQerent lipid species analyzed at 16 hpi. Results are expressed as mean ± SD of five independent replicates. Data were analysed with Student’s t-test (* p < 0.05, ** p < 0.01, *** p < 0.001; **** p < 0.0001) comparing gene expression and lipid content in uninfected and infected samples within each time point of infection.

RT-qPCR analyses of HSV-1-infected cells performed at different time points (4, 8, 16, 24, 48 hpi) showed a 4-fold increase of AGPS expression at 16, 24 and 48 hpi, when compared to uninfected sample (Figure 3b). In contrast, GNPAT expression resulted decreased by 2-fold starting from 16 hpi (Figure 3c). Additionally, FAR1, that controls the levels of fatty alcohol for plasmalogen production, did not show any significant change in gene expression between infected and uninfected cells (Figure 3d). Instead, the expression of EPT1, which is involved in the maintenance of plasmalogen levels^25^, was 3.5- and 4-fold upregulated at 16 and 24 hpi, respectively (Figure 3e).

To determine whether the infection increases plasmalogen production, a lipidomic analysis was performed on SH-SY5Y cells infected or not with HSV-1 as illustrated in Figure 3f. A 40% increase in PE-plasmalogen and PC-plasmalogen was first observed at 16 hpi with no detectable changes at earlier time-points (Figure 3g), suggesting that increase of peroxisome abundance and functionality is required at late stages of HSV-1 infection and plasmalogens, known for their essential role membrane folding, might favor the viral envelopment and release.

Notably, total PE (and PS) levels did not increase following HSV-1 infection, underscoring that the rise in PE plasmalogens is selective and not due to a general elevation of PE species. Conversely, PC levels, as well as PC-plasmalogen species, showed concurrent increases (Figure 3g, h). Moreover, lipidomic profiling also revealed a significant increase in ceramide (Cer, 2.5-fold) and sphingomyelin (SM) content (Figure 3g, h), consistent with previous reports of HSV-1-induced sphingolipid remodeling^26^.

Collectively, these results demonstrate that HSV-1 infection induces a coordinated remodeling of peroxisome-associated plasmalogen lipid production. Furthermore, several other sphingolipid species, including ceramides and sphingomyelin, were also found to be altered, indicating a broader remodeling of cellular lipid metabolism during HSV-1 infection, which will be the focus of future investigations.

### Plasmalogen biosynthesis supports HSV-1 infection in SH-SY5Y

Having established that peroxisomes are involved in and contribute to HSV-1 replication, and that plasmalogen levels are increased during herpetic infection, we next sought to determine whether plasmalogen upregulation was merely a consequence of increased peroxisome abundance or, conversely, whether peroxisome expansion reflected an increased requirement for plasmalogen biosynthesis during HSV-1 infection.

To address this question, we generated SH-SY5Y cells lacking AGPS, a key peroxisomal enzyme required for plasmalogen biosynthesis. Successful AGPS knockout was confirmed by Western blot analysis and by sequencing of the obtained clones (Figure 4a, b). Wild-type and AGPS knockout SH-SY5Y cells were then infected with the HSV-1 *McIntyre* strain at an MOI of 0.5 and analyzed at 24 hours post-infection by immunofluorescence and viral yield assay (Figure 4c).

**Figure 4.**
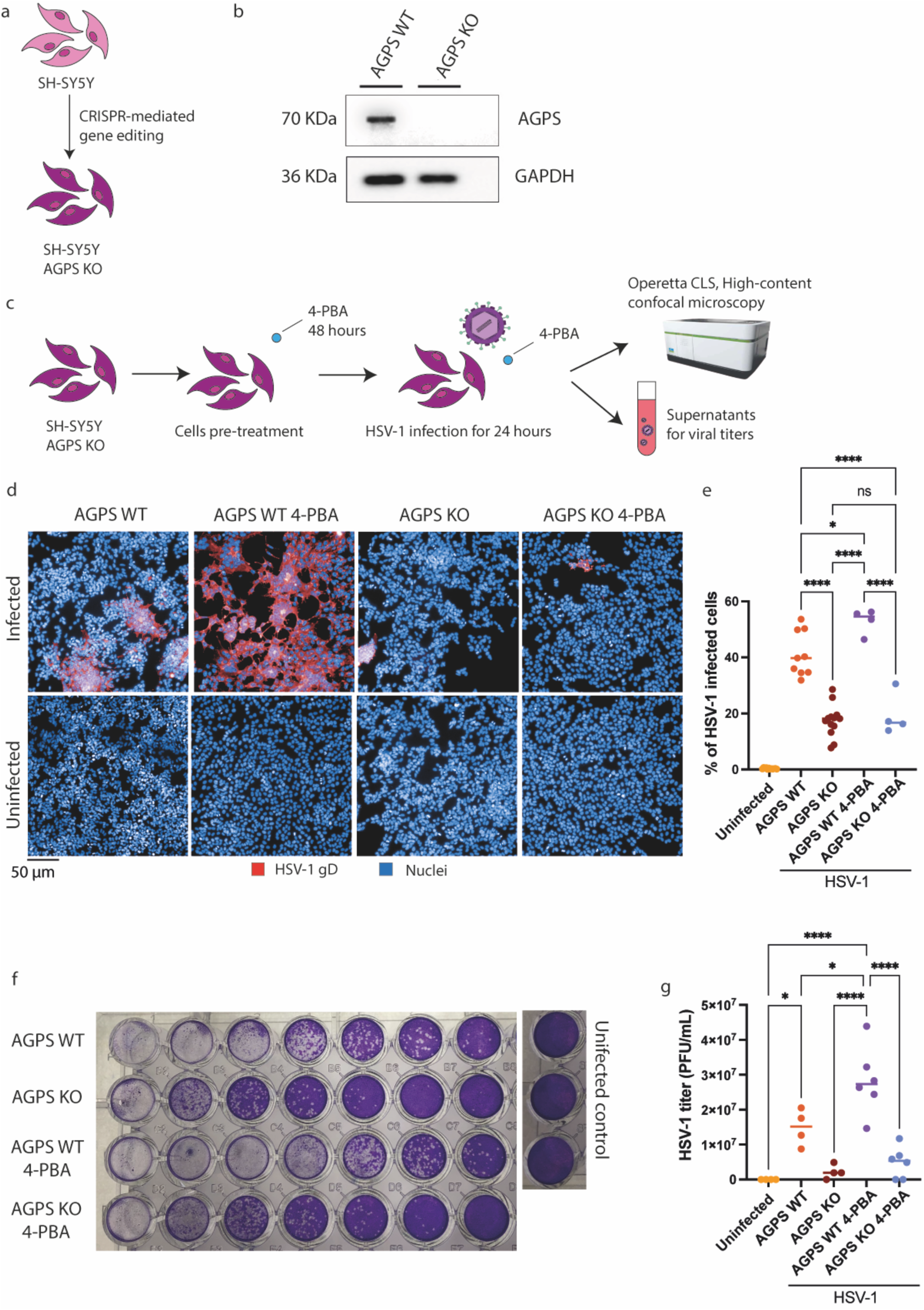
Plasmalogen production is essential for HSV-1 replication. a. Schematic representation of AGPS KO cell generation. b. Western blot image showing the AGPS KO generation. c. Schematic representation of the workflow. SH-SY5Y cells were pretreated or not with 4-PBA for 48 hours and with NCC 55-0936 for 24 hours. Then, the cells were infected with HSV-1 strain McIntyre at an MOI of 0.1 and analyzed using high content confocal microscopy and viral yield d. Panel of confocal microscopy images representing the diQerent conditions. HSV-1 using HSV-1 gD (red) and nuclei using DAPI (blue). Images were acquired using a 20x magnification. Each data point represents the mean value obtained analysing > 8×10^3^ cells. e. Analysis of the % of infected cells among the diQerent conditions: SH-SY5Y WT, AGPS KO treated or not with 4-PBA. f. Representative images of viral yield assay plate for each analyzed condition. g. Quantification of HSV-1 viral titer expressed as PFU/mL in the diQerent samples. Each data point represents a single experiment.

Both immunofluorescence analysis and viral titration showed that AGPS knockout cells displayed a marked reduction in HSV-1 infection compared with wild-type cells, with an approximately 50% decrease in infection levels. These data indicate that impairment of peroxisomal plasmalogen biosynthesis significantly compromises HSV-1 replication in neuronal-like cells.

To further investigate whether the pro-viral effect of peroxisome upregulation was specifically linked to plasmalogen production or involved additional peroxisomal functions, cells were pre-treated with 3μM 4-PBA for 48 hours before the infection. In wild-type cells, 4-PBA-mediated peroxisome upregulation enhanced HSV-1 infection. In contrast, this effect was lost in AGPS knockout cells, which were unable to synthesize plasmalogens efficiently (Figure 4d-g). Thus, increasing peroxisome abundance promoted HSV-1 infection only when plasmalogen biosynthesis was intact.

Overall, these findings suggest that the HSV-1-induced increase in peroxisomes is functionally linked to the viral requirement for plasmalogen biosynthesis. Rather than representing a passive consequence of peroxisome expansion, plasmalogen upregulation appears to be an active metabolic adaptation exploited by HSV-1 to support efficient infection and viral production.

### HSV-1 infection increases peroxisomal density in hiPSC-derived neurons

Given the pronounced effects of HSV-1 infection on peroxisomes in SH-SY5Y cells, we investigated whether similar changes occur in a human neuronal model derived from hiPSC-derived neural progenitor cells (NPCs). To do so, mature neurons were first obtained by differentiating NPCs for 30 days and then infected with the neurotropic HSV-1 *17syn+* strain at an MOI of 1 and analyzed at multiple time points (Figure 5a).

**Figure 5.**
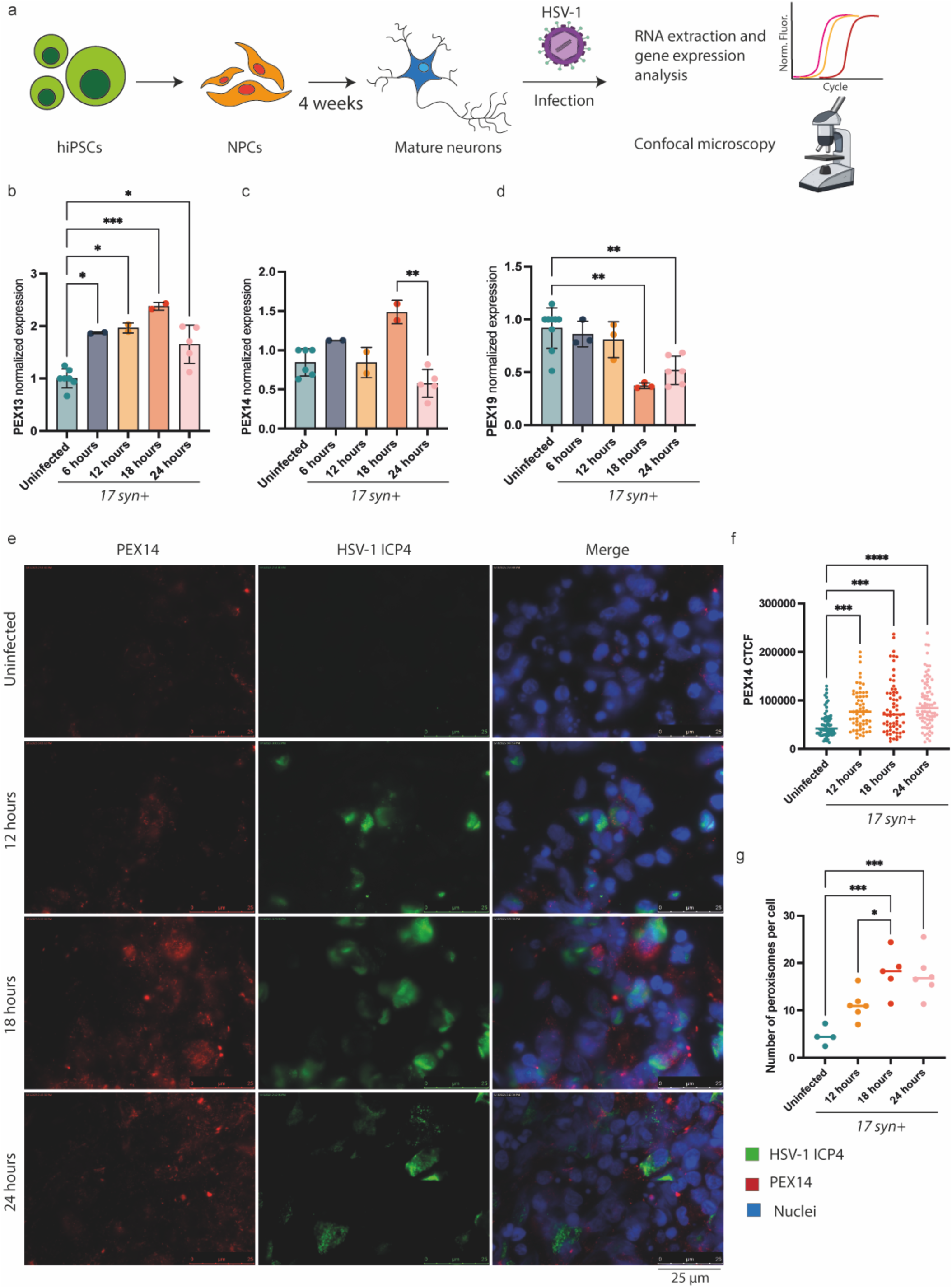
HSV-1 infection increases peroxisomes density in hiPSC-derived neurons. a. Experimental workflow. NPCs were generated from hiPSCs, diQerentiated into mature neurons for 30 days, and then infected with HSV-1 strain 17syn+ at an MOI of 1. Gene expression and peroxisome quantification were performed at 6, 12, 18, and 24 hpi by RT-qPCR and confocal microscopy respectively. Results are expressed as mean ± SD from independent biological replicates. b-d. Gene expression analyses of PEX13, PEX14, and PEX19 at indicated time point post-infection. e. Representative confocal microscopy images showing peroxisomes (PEX14, red), HSV-1 (ICP4, green), and nuclei (Hoechst, blue). Images were acquired at 63x magnification. f. Quantification of PEX14 fluorescence intensity at 12, 18, and 24 hpi relative to uninfected controls. Each data point represents the mean of PEX14 fluorescence in a defined image area normalized on the selected area background. g. Quantification of the number of peroxisomes (AU) per cell at 12, 18, and 24 hpi compared with uninfected neurons. Each data point represents the mean value obtained analysing >35 cells. Results are expressed as mean ± SD from six independent biological replicates. Statistical significance was determined by one-way ANOVA (* p < 0.05, ** p < 0.01, *** p < 0.001; **** p < 0.0001; α=0.5).

To determine whether peroxisomal alterations were directly linked to viral infection rather than being an artefactual consequence of the tumor-derived phenotype of SH-SY5Y cells, we examined PEX gene expression by qRT-PCR, and PEX14 fluorescence intensity and peroxisome number by immunocytochemistry in the presence or not of HSV-1 infection. PEX13 transcript levels doubled as early as 6 hpi (Figure 5b), whereas PEX14 expression showed a non-significant increase at 18 hpi (Figure 5c). PEX19 expression remained stable at early time points but declined after 18 hpi (Figure 5d).

Despite limited transcriptional changes, immunofluorescence analysis, revealed a 55% increase in PEX14 signal at 12 hpi, which persisted at significantly higher levels compared to uninfected controls through 24 hpi (Figure 5e, f). Since PEX14 mRNA levels did not significantly change, this effect likely reflects reduced protein degradation rather than increased gene transcription.

Quantification of peroxisome number per cell confirmed a significant increase in peroxisomal density, from the average of 4.4 peroxisomes per cell (AU) in uninfected neurons to 18.3 at 18 hpi (Figure 5g). Representative confocal images of the analyzed time points are shown in Figure 5e.

Together, these findings are consistent with results in SH-SY5Y cells and demonstrate that HSV-1 infection directly induces peroxisomal proliferation in human neurons, independent of the cellular model employed.

### Peroxisomal lipid metabolism supports HSV-1 replication in hiPSC-derived neurons

Given that HSV-1 infection promoted peroxisome proliferation and increased plasmalogen biosynthesis in SH-SY5Y cells, we next investigated how viral infection remodels the lipidome of mature hiPSC-derived neurons, a model that more closely reflects the metabolic environment of human cortical neurons. We focused our attention on plasmalogen lipids (pPE). Cells were infected with HSV-1 strains KOS or *17syn+* (MOI = 1) and analyzed at multiple time points post-infection (Figure 6a).

**Figure 6.**
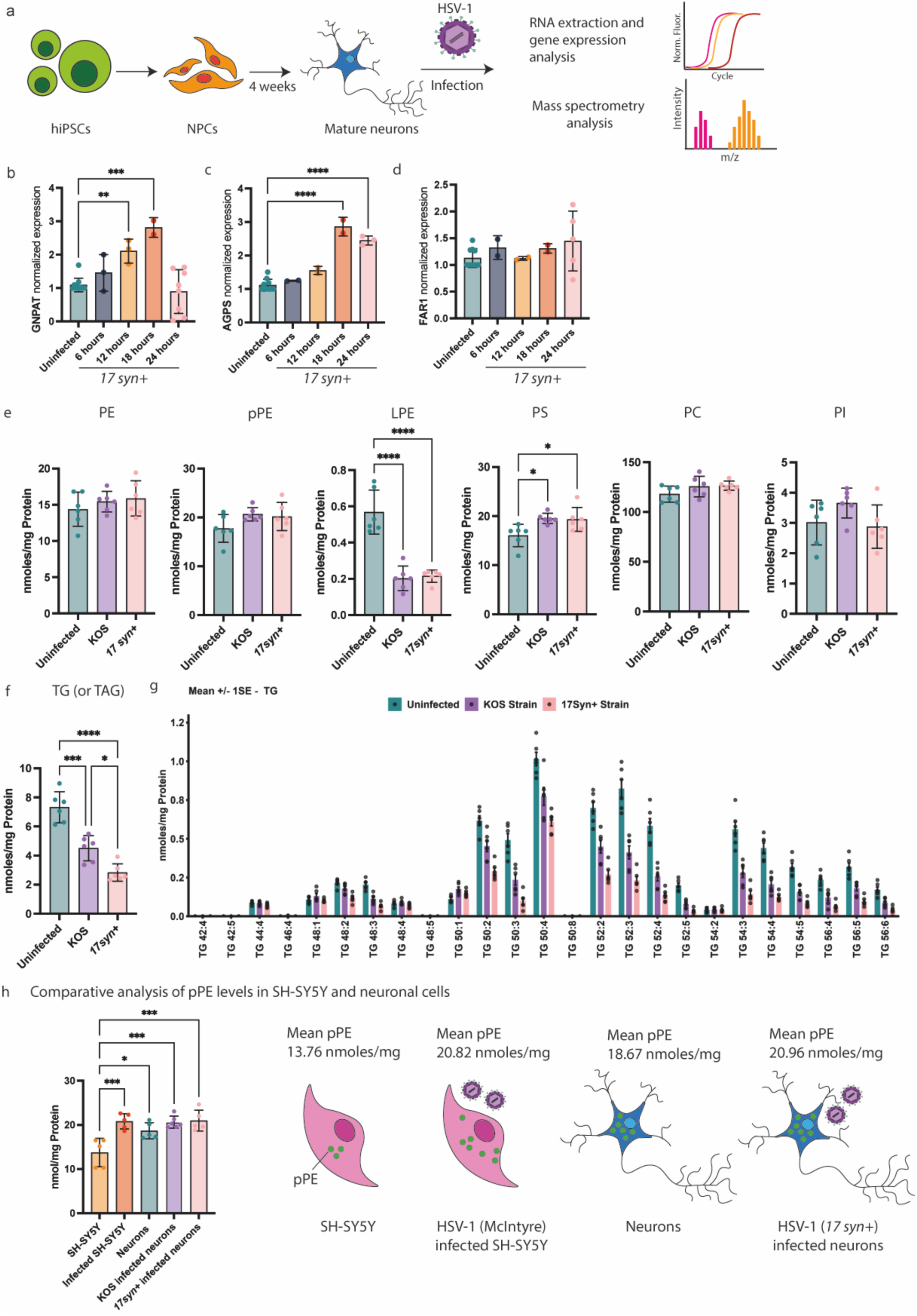
HSV-1 increases plasmalogen lipids in mature neurons as it does in SH-SY5Y cells. a. Schematic representation of the experimental workflow. Briefly, NPCs were diQerentiated for 4 weeks and infected with HSV-1 strain 17syn+ and KOS (1 MOI). Then, gene expression and lipidomic analyses were performed. b-d. Gene expression analyses of plasmalogen biosynthetic enzymes (GNPAT, AGPS, FAR1). e. Panel of absolute quantifications (nmoles/μg of protein) of diQerent lipid species in uninfected and KOS- and 17syn+-infected neurons. f. TAG absolute quantification in uninfected compared to infected counterparts. g. Graph showing the levels of TAG species. TAG with long and unsaturated chains are mostly downregulated in infected samples. h. Comparative analysis of absolute quantities of pPE in SH-SY5Y and mature neurons with or without HSV-1 infection, with a graphical representation (left panel). Results are expressed as mean ± SD from independent four or five biological replicates. Statistical significance was determined by one-way ANOVA (* p < 0.05, ** p < 0.01, *** p < 0.001; **** p < 0.0001).

To assess whether ether-lipid biosynthetic genes responded to infection, we measured the gene expression of GNPAT, AGPS, and FAR1.

GNPAT expression rose 1.5-fold at 6 hpi, 2-fold at 12 hpi, and peaked at ∼2.5-fold at 18 hpi before returning to baseline at 24 hpi (Figure 6b). AGPS was significantly upregulated (∼2.5-fold) at 18–24 hpi for the *17syn+* strain, whereas FAR1 expression remained unchanged throughout infection (Figure 6c, d).

Comprehensive shotgun lipidomics revealed that the most significant alterations involved a reduction in triacylglycerols (TAGs) and lysophosphatidylethanolamine (LPE) species. Quantitatively, TAG content decreased by ∼1.6-fold with the KOS strain and by ∼2.6-fold with the more neurovirulent *17syn+* strain, indicating strain-dependent lipid utilization (Figure 6e, f). This depletion was driven primarily by long-chain TAGs containing very-long-chain fatty acids (VLCFAs; C20–C24), while short-chain TAGs (C14–C16) remained largely unchanged (Figure 6g). Because VLCFA β-oxidation occurs exclusively in peroxisomes^27^, these results confirm an active engagement of peroxisomal lipid metabolism during HSV-1 replication. The decrease in LPE species may reflect a diversion of ethanolamine toward enhanced membrane phospholipid turnover, possibly associated with increased plasmalogen turnover required for viral envelope formation, even if net plasmalogen levels remained stable.

Instead, unlike in undifferentiated SH-SY5Y cells, plasmalogen levels (pPE) did not significantly increase upon HSV-1 infection in hiPSC-derived mature neurons. However, the pPE quantification in SH-SY5Y cells showed a mean concentration of 13.76 nmol/mg of protein, compared with the significantly higher concentration of 18.67 nmol/mg of protein measure in hiPSC-derived neurons (Figure 6h). Given the already elevated basal levels of pPE in mature neurons, this could explain why HSV-1 is unable to further increase the cellular plasmalogen lipid content. These results, showing a physiological higher pPE level in mature neurons, are consistent with the well-described feedback regulation of ether-lipid metabolism: when plasmalogen levels are high, FAR1 protein is downregulated via accelerated degradation, thereby limiting further synthesis^6^. Therefore, in mature hiPSC-derived neurons, HSV-1 likely exploits pre-existing plasmalogen pools without inducing additional synthesis, even though the virus upregulates other enzymes in the pathway (AGPS and GNPAT).

### Peroxisome remodeling is independent of HSV-1 strains in human brain organoids

After demonstrating that peroxisomal remodeling is HSV-1-specific and occurs in both neuroblastoma-derived and primary neuronal models, we next investigated whether this effect was strain-dependent or represented a conserved feature across HSV-1 isolates.

To address this, we used human brain organoids, which provide a higher degree of cellular heterogeneity and better recapitulate the *in vivo* brain environment. Organoids were differentiated for at least 40 days and then infected for three days with two HSV-1 strains, *17syn+* and KOS. Following sectioning, samples were stained for PEX14 (peroxisomes) and ICP4 (HSV-1) and analyzed by confocal fluorescence microscopy (Figure 7a) as previously described by D’Aiuto and colleagues^16^. Both viral strains induced a robust peroxisomal response, with the average peroxisome number (AU) per cell increasing by 61% for *17syn+* and 71% for KOS compared with uninfected controls. Consistently, PEX14 fluorescence intensity was increased by 61% and 67% for *17syn+*- and KOS-infected organoids, respectively (Figure 7b-h). These findings demonstrate that HSV-1–induced peroxisome proliferation is a conserved phenomenon, independent of viral strain.

**Figure 7.**
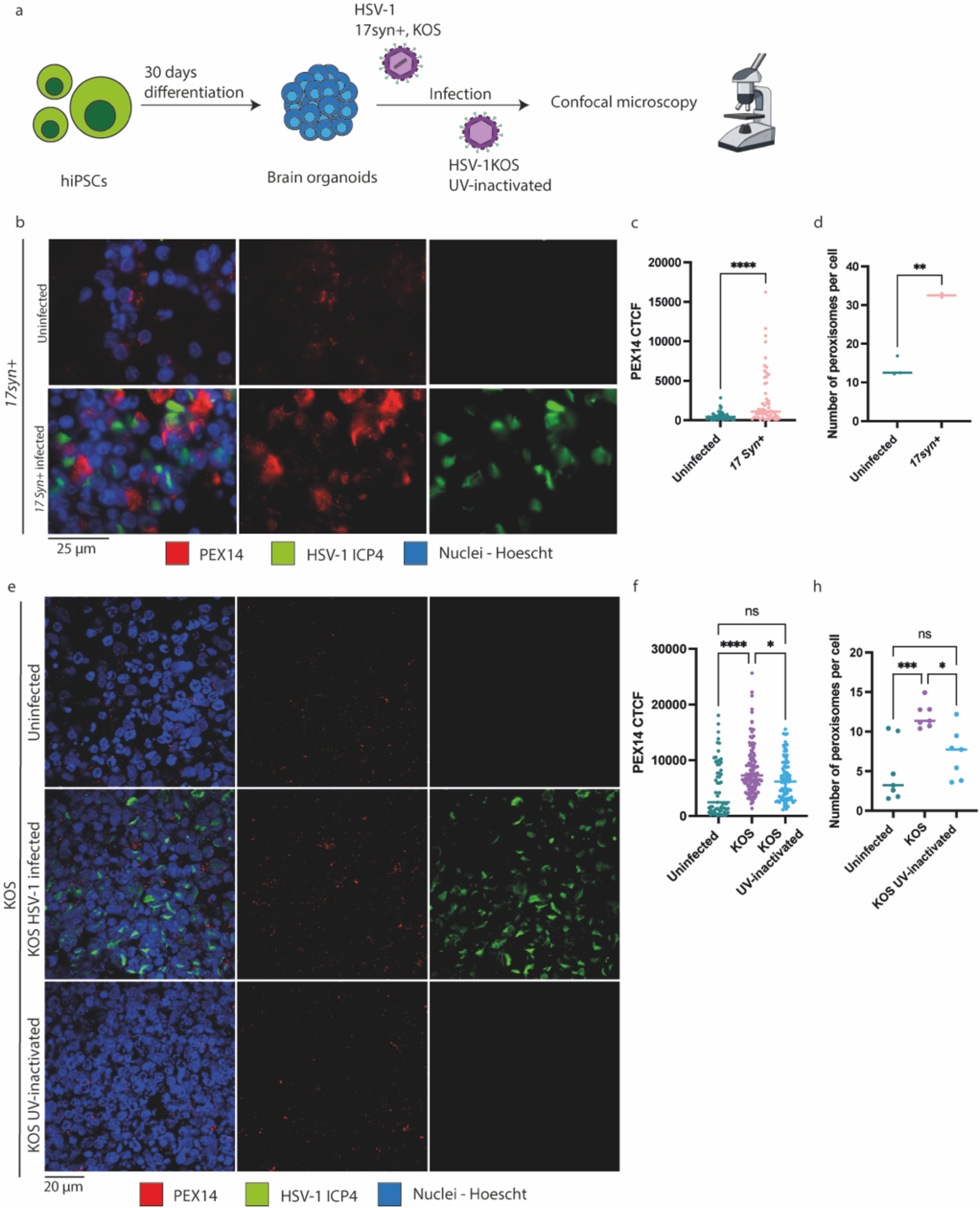
HSV-1 infection increases peroxisome abundance in human brain organoids. a. Experimental workflow. Brain organoids were generated from hiPSCs, matured for ≥40 days, and infected with HSV-1 strain 17 syn+, KOS, or UV-inactivated KOS (3000 PFU per organoid) for 3 days. Sample were then processed for and analyzed by confocal microscopy. b. Representative confocal images of HSV-1 17 syn+-infected organoids stained for peroxisomes (PEX14, red), HSV-1 (ICP4, green), and nuclei (Hoechst, blue). Images were acquired at 63x magnification. c. Quantification of PEX14 fluorescence intensity in 17syn+–infected organoids compared with uninfected controls. Each data point represents the mean of PEX14 fluorescence in a defined image area normalized on the selected area background. d. Quantification of peroxisome number (AU) per cell in in 17syn+–infected organoids relative to controls. Each data point represents the mean value obtained analysing >35 cells. e. Representative confocal microscopy images of KOS-infected, and UV-inactivated KOS-infected organoids stained as above. Images were acquired at 63x magnification. f. Quantification of PEX14 fluorescence intensity in KOS- and UV-inactivated KOS-infected organoids relative to controls. Each data point represents the mean of PEX14 fluorescence in a defined image area normalized on the selected area background. h. Quantification of peroxisome number per cell in KOS, UV-inactivated KOS, and control organoids. Each data point represents the mean value obtained analysing >35 cells. Results are expressed as mean ± SD from independent biological replicates. Statistical significance was determined by Student’s t-test or one-way ANOVA (* p < 0.05, ** p < 0.01, *** p < 0.001; **** p < 0.0001).

Strikingly, infection with UV-inactivated HSV-1 KOS, which can enter cells but is unable to replicate due to DNA damage, did not trigger a substantial reorganization of the peroxisomal compartment, excluding a generic antiviral response in inducing the peroxisomal remodeling. Hence, these data indicate that peroxisomal activation in brain organoids is a general feature of HSV-1 infection, independent of the viral strain.

### HSV-1 reactivation from latency induces peroxisomal remodeling similar to acute infection in brain organoids

To explore the potential contribution of HSV-1 reactivation to neurodegenerative diseases mechanisms through changes in peroxisomal dynamics, we employed human brain organoids as a physiologically relevant model. To recapitulate the *in vivo* neuronal environment where HSV-1 establishes latency and undergoes recurrent reactivation, we generated a latency model by treating brain organoids with a combination of 5BVdU and IFNα prior to and during infection with HSV-1 (strain KOS, 1500 PFU) for 15 days in the continued presence of antivirals. Reactivation was subsequently induced by the administration of either NaB or PI3Ki, and samples were then stained for PEX14 (peroxisomes) and ICP4 (HSV-1). Then, organoids were analyzed by confocal microscopy, as schematically illustrated in Figure 8a. The validation of the latency induction protocol and subsequent reactivation was performed following D’Aiuto et al.^16^.

**Figure 8.**
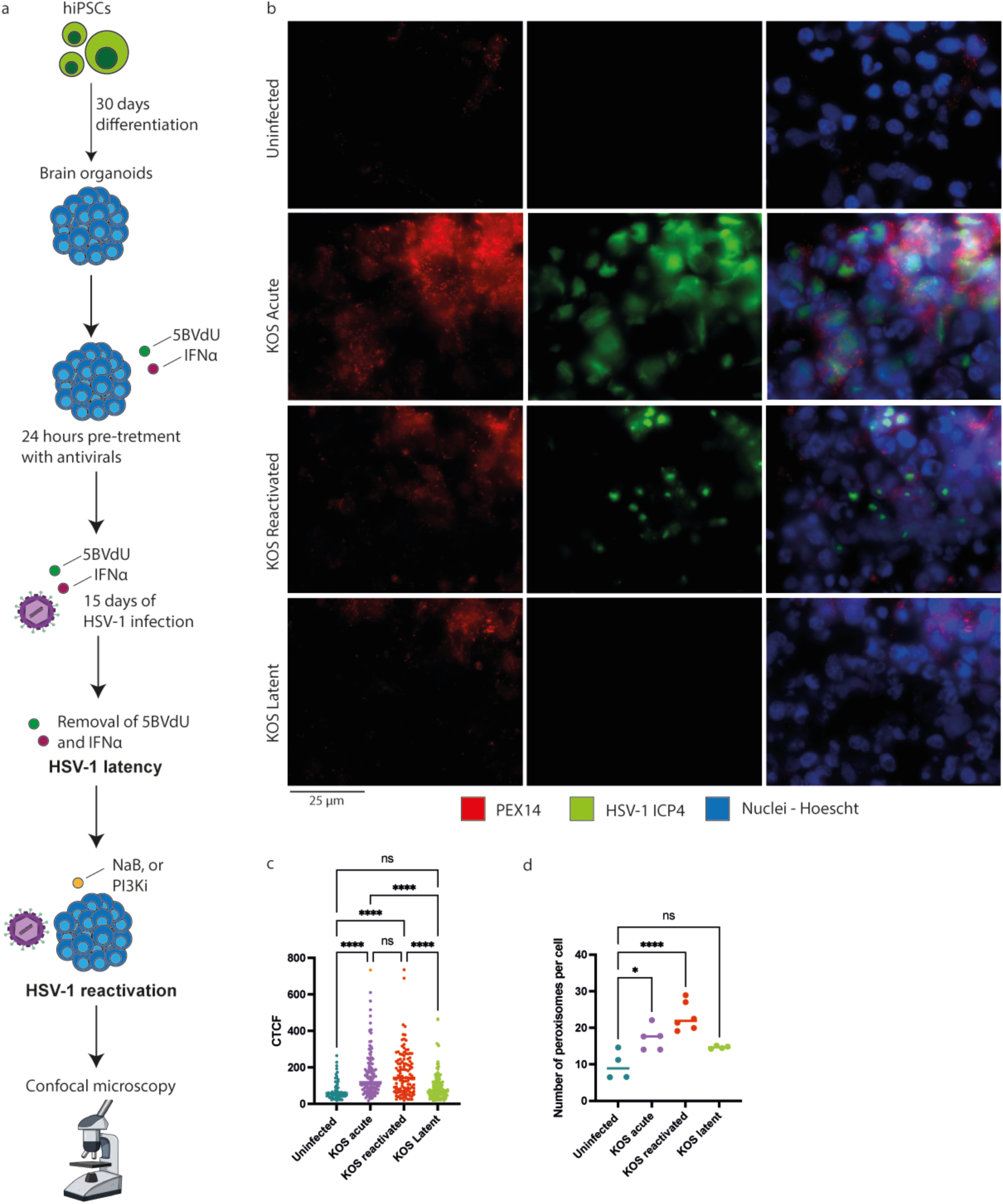
Reactivation from latency induced a rearrangement of peroxisomal compartment. a. Schematic representation of the experimental workflow. hiPSCs were used to form cortical organoids. Organoids were then pretreated with antivirals (5BVdU and IFNα) and infected with 1500 PFU of HSV-1 strain KOS. The induction of viral reactivation was then performed using NaB or PI3Ki. Samples were analyzed using confocal microscopy. b. Representative 63x magnification images acquired at confocal microscopy. The samples were then stained for peroxisomes (PEX14, red), HSV-1 (ICP4, green), nuclei (Hoechst, blue). c. Analysis of normalized fluorescence intensity of PEX14 in uninfected, and KOS infected at diQerent stages of the viral life cycle. d. Analysis of the number of peroxisomes (AU) per cell. The analysis of the number of peroxisomes per cells was performed after an additional acquisition using Operetta CLS high content confocal microscopy and the Harmony Software for image analysis. Each data point represents the mean value obtained analysing >200 cells. Results are expressed as mean ± SD of independent replicates. Data were analysed with One-Way ANOVA (* p < 0.05, ** p < 0.01, *** p < 0.001; **** p < 0.0001).

Quantitative image analyses revealed that HSV-1 reactivation led to a robust ∼54 % increase in peroxisomal fluorescence intensity, comparable to that observed during acute infection. In contrast, latently infected organoids showed no peroxisomal increase, remaining similar to uninfected controls (Figure 8b, c). Consistent results were obtained when quantifying the number of peroxisomes per cell (Figure 8d).

These findings demonstrate that peroxisomal remodeling occurs during HSV-1 reactivation, but not during latency. This evidence suggests that recurrent HSV-1 reactivation events may sustain peroxisomal reorganization and potentially impair peroxisome-associated neuronal functions, thereby contributing to neurodegenerative disease mechanisms.

## Discussion

This study identifies peroxisomes as dynamic and functionally relevant organelles during HSV-1 life cycle in human neurons. By combining tumor-derived neuroblastoma cells, hiPSC-derived neurons, and 3D brain organoids, we demonstrate that HSV-1 profoundly remodels the peroxisomal compartment, coupling organelle biogenesis with metabolic changings. These results uncover a new aspect of virus-host interaction in the nervous system, linking HSV-1 infection to peroxisomal and related lipid metabolism remodeling and providing insights on how recurrent HSV-1 reactivation may affect neuronal health.

Peroxisomes are increasingly recognized as key regulators of neuronal homeostasis and lipid metabolism, particularly in the biosynthesis of plasmalogens, essential membrane lipids supporting vesicle formation and fusion, myelination, and antioxidant defense^28–31^. Indeed, altered peroxisomal activity and plasmalogen imbalance have been associated with Alzheimer’s and other neurodegenerative disorders, highlighting their relevance to neuronal integrity^5,12^. Our findings expand their role by showing that HSV-1 actively induces peroxisome biogenesis across different neuronal models. Transcriptional upregulation of PEX13, PEX14, and PEX19, together with increased peroxisome number and PEX14 fluorescence signal, indicate that HSV-1 stimulates peroxisome formation in both SH-SY5Y cells, hiPSC-derived neurons and brain organoids. Pharmacological modulation of peroxisomes further supports this view: enhancing peroxisome proliferation with 4-PBA promoted viral replication, whereas inhibition with NCC 55-0396 markedly reduced infection efficiency. These results demonstrate that peroxisome biogenesis is not a host defense mechanism but rather a process exploited by HSV-1 to sustain its replication cycle. In line with this, UV-inactivated HSV-1 in brain organoids did not retain the ability to increase peroxisome abundance, further confirming the active role played by the virus on these cellular organelles.

Lipidomic profiling revealed broad remodeling of host lipid metabolism following HSV-1 infection, including increases in ceramides, sphingomyelins and plasmalogens (pPE and pPC) in SH-SY5Y cells. Contrarily, in mature hiPSC-derived neurons lipidomic analysis revealed very mild and non-significant changes in pPE levels, even though the virus upregulates the transcription of two enzymes in the pathway (AGPS and GNPAT). However, comparing the pPE concentrations between SH-SY5Y and mature neurons, we observed a higher physiological level of plasmalogen lipids in neurons, equal to the ones after HSV-1 infection in SH-SY5Y. In line with this, FAR1 transcription does not show any variation during infection. This may suggest a possible activation of the regulatory feedback of ether-lipid metabolism: when plasmalogen levels are high, FAR1 protein is downregulated via accelerated degradation, thereby limiting further synthesis^32,33^. Although this observation still requires validation in hiPSC-derived neurons, it suggests that mature neurons possess a limited capacity to further enhance plasmalogen biosynthesis, having already high levels of pPE. Consequently, in mature hiPSC-derived neurons, HSV-1 likely exploits pre-existing plasmalogen pools without inducing additional synthesis. The alterations in our lipidomic data, both in SH-SY5Y cells and hiPSC-derived neurons, align with previous studies showing HSV-1-dependent activation of ceramide and sphingolipid pathways, which facilitates viral envelopment and egress. On the other side, the upregulation of peroxisome-derived plasmalogens or the use of pre-existing high concentrations, confirmed both at the lipid and transcriptional levels, identify peroxisomes as a key metabolic hub supporting HSV-1 replication.

The functional relevance of plasmalogen biosynthesis during HSV-1 infection was further supported by AGPS knockout experiments. Loss of AGPS, a key peroxisomal enzyme involved in ether lipid and plasmalogen synthesis, reduced HSV-1 infection by approximately 50–60% in SH-SY5Y cells. Moreover, 4-PBA-induced peroxisome proliferation enhanced infection in wild-type cells but not in AGPS knockout cells. These findings indicate that HSV-1-induced peroxisome biogenesis promotes viral replication by sustaining plasmalogen production.

Because plasmalogen are essential for neuronal function^30,34,35^, repeated viral reactivation could progressively impair neuronal lipid homeostasis and contribute to chronic dysfunction. In line with the central role of peroxisome in HSV-1 infection, hiPSC-derived neurons showed a marked reduction in TAGs, especially long-chain TAGs, following HSV-1 infection. VLCFAs are specifically controlled by peroxisomal metabolism^29,36^. Indeed, the observed depletion of long-chain TAGs suggests a shift in peroxisomal lipid flux, possibly reflecting enhanced VLCFA β-oxidation.

This hypothesis that peroxisomes and, more specifically, plasmalogen are used by HSV-1 for its own replication is sustained by the biophysical features of plasmalogens, that make them crucial for membrane architecture and vesicle fusion processes, possibly facilitating viral replication and egress from host cells, consistent with what was observed during HCMV infection^14^. Indeed, HCMV upregulates plasmalogen lipids at late stages of infection to facilitate the secondary envelope of the virus^14^.

Therefore, to explore the potential link between HSV-1 infection and neuronal health, we employed a more physiologically relevant experimental system that better recapitulates the *in vivo* infection of human neurons. Our latency-reactivation model in brain organoids demonstrates that peroxisomal remodeling is restricted to active viral replication and is absent during latency. Reactivation fully restored the increase in peroxisome number and PEX14 signal observed in acute infection, providing direct evidence that HSV-1 reactivation episodes repeatedly perturb peroxisomal homeostasis. Given that HSV-1 establishes lifelong latency in neurons, recurrent reactivation may cause cumulative metabolic stress, contributing to neurodegenerative cascades over time. The use of 3D brain organoids, which recapitulate neuronal complexity and tissue architecture, strengthens the translational relevance of these findings and bridges the gap between simplified 2D cultures and the human brain environment.

While the present data strongly support a model in which HSV-1 hijacks peroxisomal function, several open questions remain. The viral determinants responsible for triggering peroxisome biogenesis have to be identified. Moreover, it remains to be determined at which step of the HSV-1 replication cycle plasmalogens are required, and whether they mainly contribute to membrane remodeling events involved in viral assembly, envelopment, or egress.

## Acknowledgements

This project was funded by ***Progetto PNRR*** – Spoke 3 “Neurodegeneration” – Sviluppo di terapia genica e farmaci con tecnologia a RNA, ***Istituto Superiore di Sanità*** ‘Decoding the pathogenesis of SARS-CoV-2: Investigating host and viral biomarkers in vitro and in an animal model across multiple viral variants’ - RIPREI2023_f5b001ceb717. ***National Research Programme and for Projects of National Interest (NRP)***: Harnessing the host innate immune response against single stranded RNA-positive emerging viruses. Prot.P20222HHXA, the ***Boehringer Ingelheim Fonds Travel Grant*** and the ***Blanceflor Foundation Short-term Scholarship***. We acknowledge CISUP—Centre for Instrumentation Sharing—University of Pisa for the use of Operetta CLS imaging facility

## Authorship contribution statement

**Conceptualization:** M.L, C.F., R.A, **Writing and Data Curation:** C.F., M.L, R.A. **Methodology and Data analysis:** A.D.C, I.G, C.P, E.I, F.F, G.S, D.F**. Investigation, review:** M.A.W, L.D.A. G.F. **Supervision, Funding acquisition and review**: M.L, M.P. **Visualization:** C.F.

