## Supplementary materials for "Peroxisome dynamics during HSV-1 life cycle in human neurons"

*Table 1. Media composition for neuronal maintenance and differentiation into mature neurons and 3D cortical organoids.*

| Medium | Composition |
|---|---|
| **NP expansion medium** (Gibco, Thermo Fisher Scientific, USA) | DMEM/F12 supplemented with 0.5% N2, 1 mM L-glutamine, 1% non-essential amino acids, 50U/mL penicillin G, 50 mg/mL streptomycin, and 0.1 mM 2-mercaptoethanol |
| **Incomplete Neurobasal Medium** (Gibco, Thermo Fisher Scientific, USA) | Neurobasal medium, 1 mM L-glutamine, 50 U/mL penicillin G, 50 mg/mL streptomycin, 2% B27 supplement, 10 ng/mL brain-derived neurotrophic factor (BDNF), 3 mM CHIR9902110, 1 mM Dorsomorphin, 10 mM Forskolin, and 10 mM Rock inhibitor |
| **Complete Neurobasal Medium** (Gibco, Thermo Fisher Scientific, USA) | Neurobasal medium, 1 mM L-glutamine, 50 U/mL penicillin G, 50 mg/mL streptomycin, 2% B27 supplement, and 10 ng/mL BDNF |
| **Neuronal Medium** | DMEM/F12 supplemented with 0.5% N2, 1 mM L-glutamine, 1% non-essential amino acids, 2% B27, Heparin, 20 µg/mL rhFGF and rhEGF |
| **CODMI Medium** | DMEM/F12 combined in a ratio of 1:1 with Neurobasal medium, supplemented with 0.5% N2, 1mM L-glutamine, 1% non-essential amino acids, 50U/mL penicillin G, 50 mg/mL streptomycin, 2% B27, 20 µg/mL rhFGF and rhEGF |
| **CODMII Medium** | 1:1 combination of DMEM/F12 and Neurobasal Medium, supplemented with 0.5% N2, 1mM L-glutamine, 1% non-essential amino acids, 50U/mL penicillin G, 50 mg/mL streptomycin, 2% B27, and 10 µg/mL BDNF. |

*Table 2. RT-qPCR primer pairs used for the analysis.*

| Target | Forward primer | Reverse primer |
|---|---|---|
| β-actin | AAGGAGAAGCTGTGCTACGTC | AGACAGCACTGTGTTGGCGTA |
| HPRT | CATTATGCTGAGGATTTGGAAAGG | CTTGAGCACACAGAGGGCTACA |
| PEX13 | ACCGAGCAGCTACCTCAGCAAA | TCATCCTCACCACTTGCCCAGT |
| PEX14 | GAGAGGACAGAAAGCAGCTGGA | AAGCTCCTGGATCTTCTGCTGC |
| PEX19 | CCTACTCTCCAAGGATGTGCTG | ATGACGCTGTGCTGCTCCTGAT |
| FAR1 | GTGGTCTCTTTATTGCGGCAGG | AATACCAGGCTGCCGCAAGACT |
| GNPAT | CACAAACTGCGTCTTGGAGCCA | GCTCCTGGATGGCTCTTTGTAG |
| AGPS | GGCTGGCATAACAGGACAAGAG | CTTCATGCCTGATGCGCGAGTA |
| EPT1 | GTGGATACCAATCCACTTTCTCTG | GAATACGACCAGCAGAAAGCCAG |
