## Supplementary figures for "Peroxisome dynamics during HSV-1 life cycle in human neurons"

Supplementary figure 1 (S1): 4-PBA and NCC 55-0396 treatment toxicity on SH-SY5Y cells.

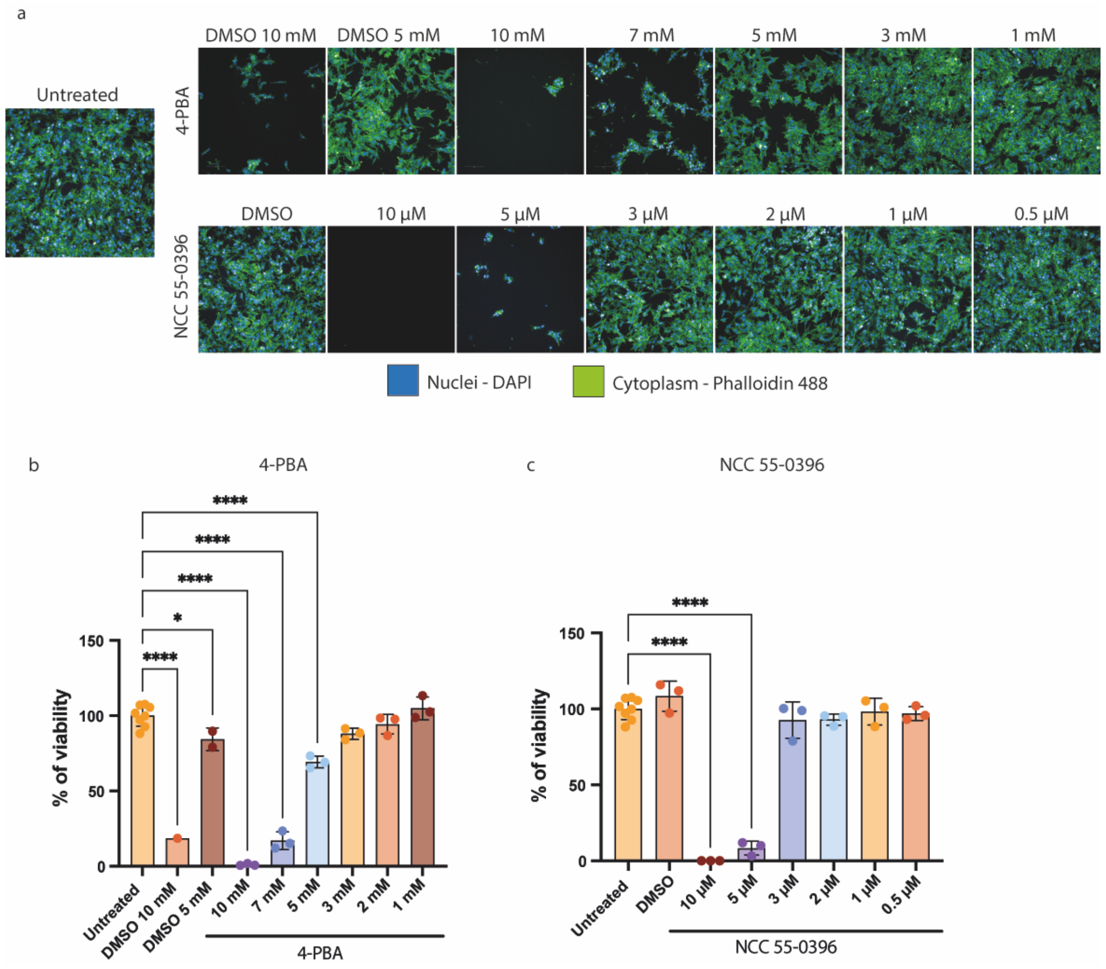


***Supplementary figure 1**. **Toxicity of 4-PBA and NCC 55-0396 treatment of SH-SY5Y cells**. a. Representative 20x magnification microscopy images of SH-SY5Y treated with 4-PBA and NCC 55-0396 at different concentrations and relative controls. Nuclei were labelled using DAPI (blue) and cytoskeleton using phalloidin (green) b. Percentage of cell viability following treatments with different concentration of 4-PBA for 48 hours. The cell viability was calculated counting the number of nuclei per sample, considering our untreated control as the 100% of cell viability. c. Percentage of cell viability of cells treated with different concentration of NCC 55-0396 for 24 hours. The cell viability was calculated as above. Results are expressed as mean ± SD of three independent replicates. Data were analysed with One-Way ANOVA (* $p < 0.05$, ** $p < 0.01$, *** $p < 0.001$; **** $p < 0.0001$).*

Supplementary figure 2 (S2): Analysis of peroxisome number in uninfected cells in HSV-1 infected well

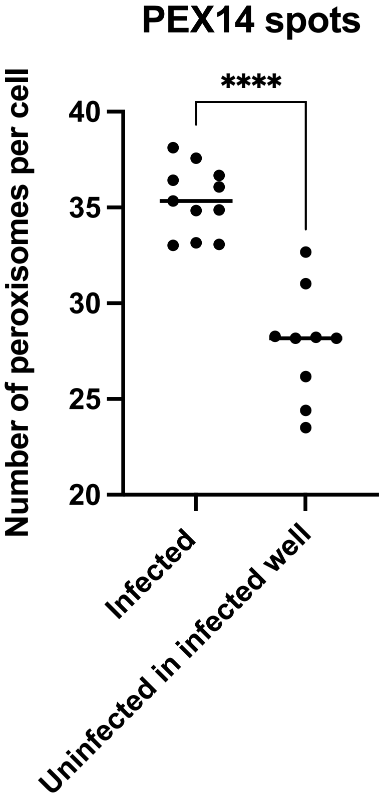


***Supplementary figure 2. Peroxisomal remodeling occurs only in presence of HSV-1***. *Analysis of the number of peroxisomes in HSV-1 (McIntyre strain) infected cells and uninfected cells in the same well. The confocal microscopy images were acquired using the high content microscopy Operetta CLS at 40x magnification. The analysis pipeline was performed using Harmony Software. Results are expressed as mean ± SD of three independent replicates. Each replicate is the mean of >6000 cells analyzed. Data were analysed with One-Way ANOVA (* p < 0.05, ** p < 0.01, *** p < 0.001; **** p < 0.0001).*

Supplementary figure 3: Effect of peroxisomal remodelling on ZIKV infection.

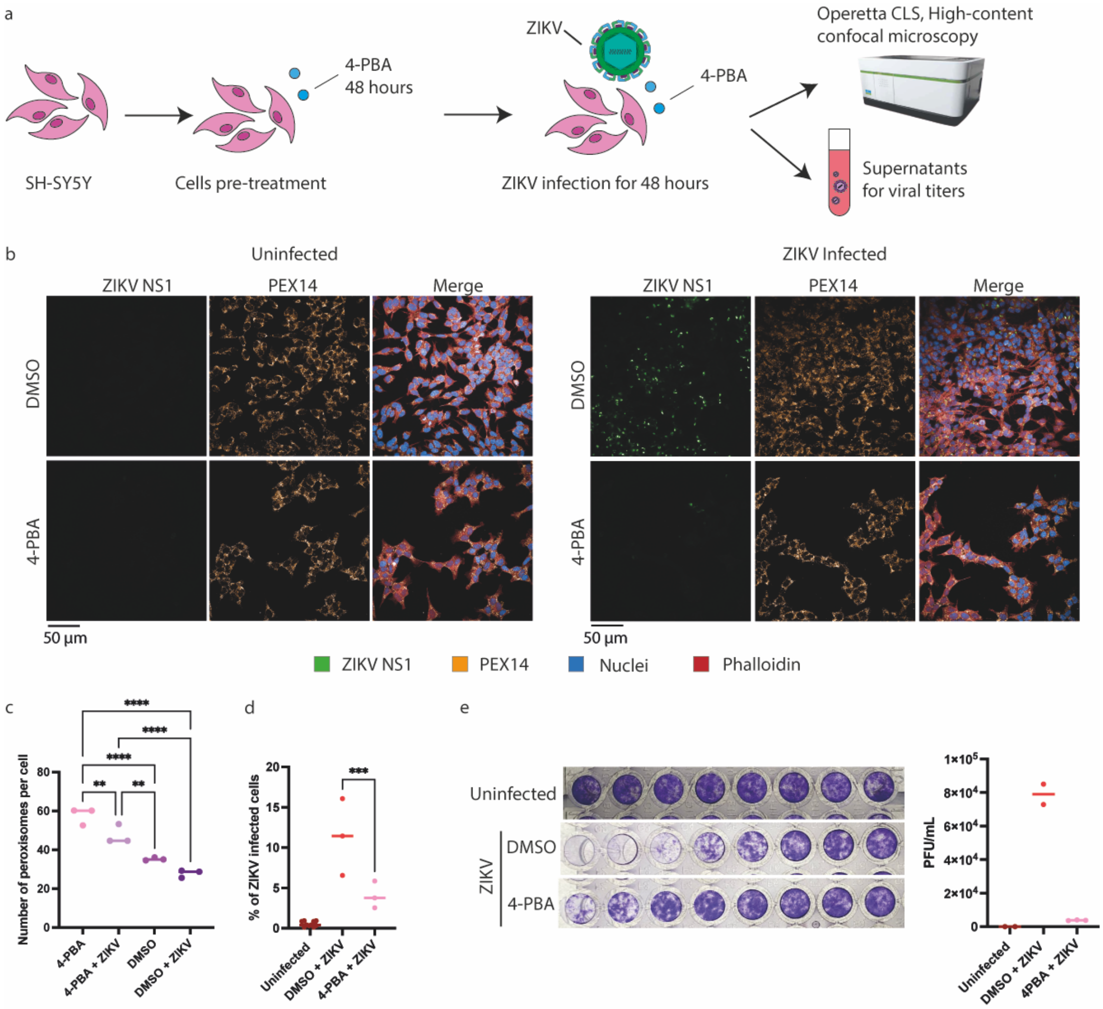


***Supplementary figure 3. 4-PBA reduces ZIKV infection***. *a. Overview of the workflow. Briefly SH-SY5Y were pretreated with 4-PBA for 48 hours. Cells were then infected with ZIKV at an MOI of 2 for 48 hours. The supernatants were collected to perform a viral yield assay, and the cells were analyzed using high content confocal microscopy. b. Panel of representative microscopy images. Peroxisomes were labelled using PEX14 (orange), ZIKV using ZIKV NS1 (green), nuclei with DAPI (blue), and cytoskeleton with phalloidin (red). The images were acquired using a 40x magnification. Each data point represents the mean value obtained analysing > 8x10$^3$ cells. c. Analysis of confocal images using Harmony software showing the number of peroxisomes per cell. d. Analysis of confocal microscopy images showing a reduction in the percentage ZIKV-positive cells compared to vehicle treated sample. e. Viral yield assay performed using supernatant of 4-PBA treated cells. The viral titer is expressed as PFU/mL. Results are expressed as mean ± SD of independent replicates. Data were analysed with One-Way ANOVA (* $p < 0.05$, ** $p < 0.01$, *** $p < 0.001$; **** $p < 0.0001$).*
